# SMILE: Mutual Information Learning for Integration of Single Cell Omics Data

**DOI:** 10.1101/2021.01.28.428619

**Authors:** Yang Xu, Priyojit Das, Rachel Patton McCord

**Affiliations:** UT-ORNL Graduate School of Genome Science and Technology, University of Tennessee, Knoxville, TN; Biochemistry & Cellular and Molecular Biology Department, University of Tennessee, Knoxville, TN

## Abstract

Deep learning approaches have empowered single-cell omics data analysis in many ways, generating new insights from complex cellular systems. As there is an increasing need for single cell omics data to be integrated across sources, types, and features of data, the challenges of integrating single-cell omics data are rising. Here, we present a deep clustering algorithm that learns discriminative representation for single-cell data via maximizing mutual information, SMILE (**S**ingle-cell **M**utual **I**nformation **Le**arning). Using a unique cell-pairing design, SMILE successfully integrates multi-source single-cell transcriptome data, removing batch effects and projecting similar cell types, even from different tissues, into the same representation space. SMILE can also integrate data from two or more modalities, such as joint profiling technologies using singlecell ATAC-seq, RNA-seq, DNA methylation, Hi-C, and ChIP data. SMILE works well even when feature types are unmatched, such as genes for RNA-seq and genome wide peaks for ATAC-seq.

## Introduction

Deep-learning-based single cell analysis has gained great attention in recent years and has been used in a range of tasks, including accurate cell-type annotation (Ma and Pellegrini 2020), expression imputation (Arisdakessian et al. 2019) and doublet identification (Bernstein et al. 2020). In these tasks, deep learning showed some striking advantages. For example, in cell-type annotation, the automatic and accurate annotation using a deep classification model saves researchers from manual cell-type annotation (Lopez et al. 2018; Kimmel and Kelley 2020; Ma and Pellegrini 2020). Another application of deep learning is data imputation and denoising. Though there has been a dramatic improvement in scRNA-seq technology, the problem of zero-inflation remains as a grand challenge in single cell transcriptomics (Lähnemann et al. 2020). Due to the difficulty of modeling technical zero values and biological zeros, deep learning has become a more appealing alternative for this task. An autoencoder (AE) is a common artificial neural network that is used to learn representations for data in an unsupervised manner. Both DeepImpute (Arisdakessian et al. 2019) and DCA (Eraslan et al. 2019) adopt a variant of AE model to impute gene expression and de-noise single cell data. These approaches and many others are revealing the power of deep learning applied to single cell genomic datasets.

Data integration is a rising challenge in single-cell analysis, as increasing numbers of single-cell sequencing datasets become available and the types of sequencing data become more diverse. Consequently, data integration becomes a key research domain for understanding a complex cellular system from different angles. However, there are few comprehensive tools for data integration to address this challenge. In single-cell transcriptomics, batch effects are usually a prominent variation when comparing data from multiple sources and removing batch effects is a critical step for revealing biologically relevant variation. Besides integrating single-cell transcriptome data, integration of multimodal single-cell data is becoming important as technological breakthroughs make it possible to capture multiple data types from the same single cell. For example, sci-CAR and SHARE-seq can simultaneously profile chromatin accessibility and gene expression for thousands of single cells (Cao et al. 2018; Ma et al. 2020). scMethyl-HiC and sn-m3C-seq can profile DNA methylation and 3D chromatin structure at the same time at single-cell resolution (Lee et al. 2019; Li et al. 2019). However, these data types do not naturally share the same feature space: transcriptomes are described using genes as features, while chromatin accessibility is reported across all intergenic spaces. Therefore, integration of multimodal data becomes more challenging. Furthermore, a new technology named Paired-Tag now achieves joint profiling of gene expression and 5 different histone marks for thousands of single nuclei (Zhu et al. 2021). This new technology introduces a further challenge of integrating more than 2 modalities. To address the challenges above in a single method, we designed a deep clustering model, SMILE, that learns discriminative representation for data integration in an unsupervised manner. In our approach, we restructured cells into pairs, and we aimed to maximize the similarity between the paired cells in the latent space. Because of this cell-pairing design, SMILE extends naturally into integration of multimodal single-cell data, where data from two sources (RNA-seq/ATAC-seq or Methyl/Hi-C) exist for each cell and thus form a natural pair. We demonstrated that SMILE can effectively project RNA-seq/ATAC-seq data, as well as Methyl/Hi-C data, into the same latent space and achieve data integration. Finally, we present a combinatorial use of SMILE models to integrate RNA-seq, H3K4me1, H3K9me3, H3K27me3, and H3K27ac data generated by Paired-Tag. SMILE performs as well or better than other methods designed for data integration while also having increased flexibility in terms of data input types.

## Results

### SMILE uses a cell pairing strategy to integrate multi-source single-cell transcriptome data

We first explain the architecture of SMILE (Fig. 1A). The main component is a multi-layer perceptron (MLP) used as an encoder that projects cells from the original feature space *X* to representation *z.* To achieve this goal, SMILE relies on maximizing mutual information between *X*and *z*. Mutual information measures the dependency of *z* on *X*. If we maximize the dependency, we can end up using low-dimension *z* to represent the high-dimension *X*. Contrastive learning is one approach to maximize mutual information, and it usually requires pairing one sample with a positive or negative sample. Then, the goal is to maximize similarity between the positive pair and dissimilarity between the negative pair in the representation *z*. Due to the lack of labels, pairing samples is a challenging task. However, treating a sample itself as its positive sample and any other cells as negative samples can be a shortcut for reframing the data into pairs, and Chen et al. demonstrated that such a simple framework can effectively learn visual representation for images (Chen et al. 2020). In single-cell data, we adopt the same framework to pair each cell in a dataset to itself. To prevent each pair from being completely the same, we add gaussian noise to each cell. Then, maximizing mutual information becomes forcing each cell to be like itself, regardless of the added noise, and dissimilar to any other cells. We find that learning a representation in which each noise-added cell is like its noise-added pair also leads to collective similarity of cells within the same cell-type (Fig. 1B-D). To implement this in the neural network, we used noise-contrastive estimation (NCE) as the core loss function to guide the neural network to learn (Wu et al. 2018). We did not directly apply NCE on representation *z*, but further reduced *z* to a 32-dimension output and *K* pseudo cell-type probabilities, by stacking two independent one-layer MLPs onto the encoder. A one-layer MLP generating a 32-dimension vector will produce rectified linear unit (ReLU) activated output, and the other will produce probabilities of pseudo cell-types with SoftMax activation. Finally, NCE was applied on the 32-dimension output and pseudo probabilities, independently. These two one-layer MLPs produce two independent outputs which both contribute to the representation learning of the encoder. Once trained, the encoder serves as a feature extractor that projects data from the original space *X* to a low-dimension representation *z*. Further implementation detail about NCE and the neural network can be found in **Methods**.

**Fig. 1.**
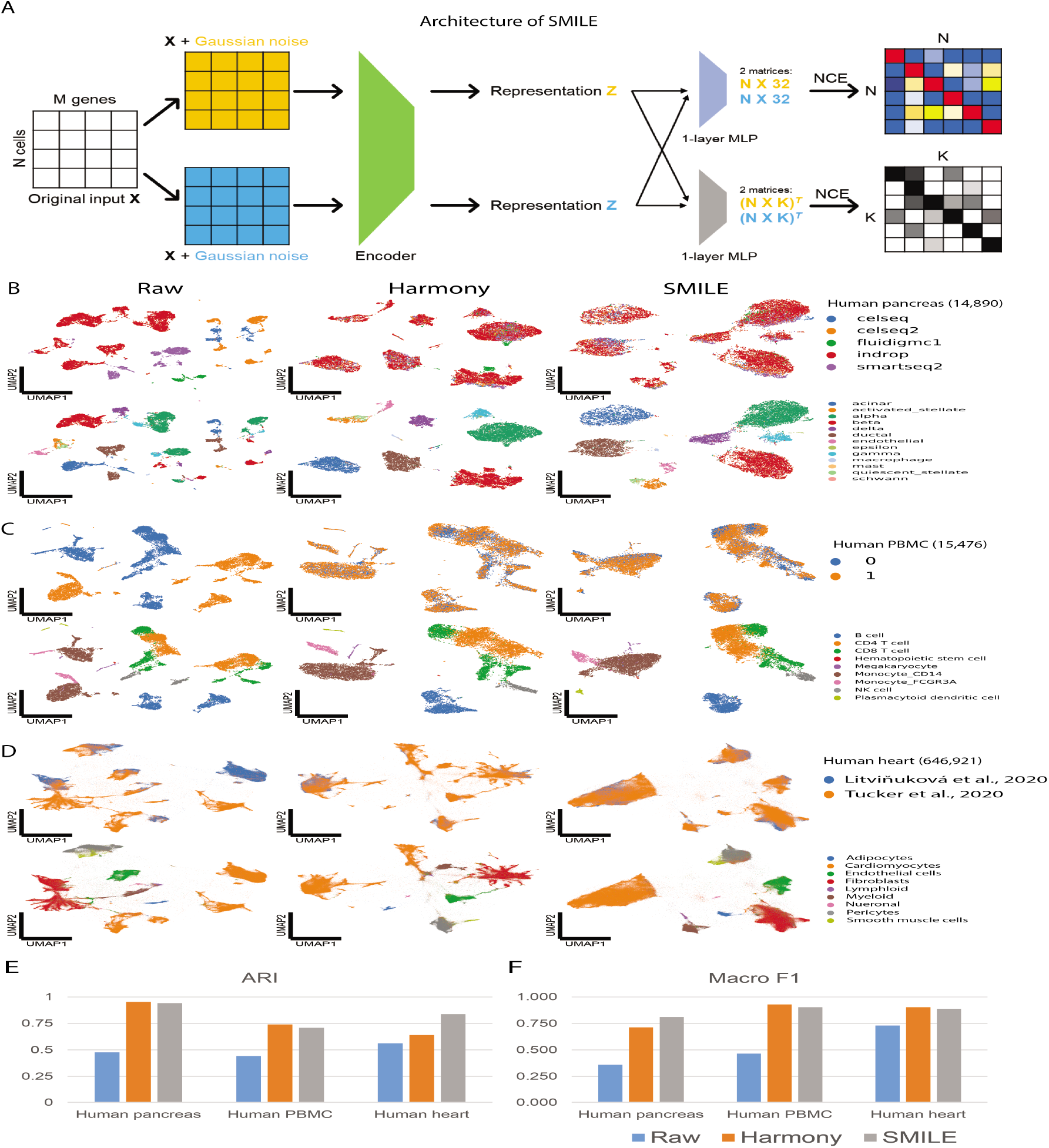
Integration of multi-source single-cell transcriptome data using SMILE. (A) Architecture of SMILE. Original input *X* represents a gene expression matrix where each row represents a cell, and each column stands for a gene. Random Gaussian noise is added to differentiate the input *X* into two *X*s, which are the same except for the added noise. Encoder (green) consists of two fully connected layers that projects *X* into 128-dimension representation *z.* Two independent fully connected layers (blue and grey) are stacked onto the encoder to further reduce *z* into 32-dimension output and *K* pseudo cell-type probabilities, respectively. Computing NCE using the 32-dimension output and *K* pseudo cell-type probabilities will generate two matrices. One is in the shape of *N* by *N (N* is the number of cells in one batch), and the other is the shape of *K* by *K*. Then gradient descending guides the model to increase values on the diagonal of the two matrices while decreasing values in the off diagonal. (B-D) UMAP visualization of integrated representations of B) human pancreas data, C) human PBMC data, and D) human heart data, using raw data, or representations learned by Harmony and SMILE. Cells in upper rows are colored according to their sources or batch ID, and cells in lower rows are colored by putative celltypes reported in original studies. (E) Evaluation of recovery of cell-types through clustering. ARI shows how well the learned representation can recover cell-types. ARI closer to 1 indicates that the clustering labels better match original cell-type labels in that study. (F) Evaluation of label transferring. SVM classifiers are trained with single source data, and then macro F1s are calculated by assigning cell types to the rest of the data sources using that classifier.

### Contrasts between SMILE and other deep learning approaches

Before applying SMILE to batch correction for multi-source single-cell transcriptome data, we present the differences between SMILE and other deep learning models that have been proposed to learn representation for single-cell data. Variants of Variational Autoencoder (VAE) models, which differ in their sampling approaches, have been proposed to learn representations for singlecell data with batch-effect correction (Lopez et al. 2018; Lotfollahi et al. 2018; Wang et al. 2019; Bahrami et al. 2020). The core component of VAE is the use of reconstruction loss, but VAE also encodes a sample in the representation where this sample is drawn from a certain distribution, for example, a Gaussian distribution (Supplementary Fig. S1A). The use of reconstruction loss does not necessarily push the representation to be discriminative, and it is also unlikely that the sampling approach based on one underlying distribution in VAE will allow samples from different classes to have different means and variances. This leads to a drawback that representations learned by VAE-based methods may not be discriminative enough for clustering. Another deep clustering approach directly models clustering loss, and two studies have demonstrated the ability to learn more discriminative representation for scRNA-seq data (Tian et al. 2019; Li et al. 2020). Both scDeepCluster and DESC require initiation of neural network through training an autoencoder (AE) model. However, the deep clustering tool CoSTA (Supplementary Fig. S1A), which we recently developed, demonstrated that clustering-loss-based deep clustering can directly learn discriminative representation without an initiation through AE (Xu and McCord 2021). The common core concept in these methods is modeling clustering loss of *K*-means, which usually would assume that samples are split uniformly among the *K* classes. However, most single cell data are not well balanced, and the number of cells belonging to dominant cell-types may far exceed cells belonging to rare but distinct cell-types. In such a case, clustering-loss-based deep clustering is likely to fail in the identification of non-dominant cell-types.

To show that the learning approach of SMILE can overcome issues above, we applied SMILE, VAE and CoSTA to single-source human pancreas scRNA-seq data (Baron et al. 2016). The number of cells in each cell-type in this dataset is not equal. 4 major cell-types (alpha, beta, gamma, and delta) range from 266 to 2507 cells, and 4 minor cell-types (macrophage, mast cell, epsilon cell and Schwann cell) are represented by as low as 13 cells. For fair comparison, we used an encoder with the same structure in all of SMILE, VAE and CoSTA and trained these 3 models in the same manner. We evaluated how clustering-friendly representations learned by these three models are by adjusted rand index (ARI) and normalized mutual information (NMI), and we trained SVM classifiers to evaluate how well minor cell types are recovered by these representations. Indeed, the representation by VAE is not clustering-friendly (Supplementary Fig. S1B). This supports our argument that VAE is less able to learn discriminative features if the underlying data layout is not inherently class discriminative. In contrast, CoSTA lacks the ability to recover non-dominant cell types, as shown by the micro F1 score and the F1 scores of each cell-type (Supplementary Fig. S1B). SMILE outperforms both VAE and CoSTA in aspects of clustering-friendly representation and recovery of minor cell types. While the three methods do not differ much in their ability to classify the 4 major cell types, SMILE shows superiority for classifying minor cell types.

### SMILE accommodates many single cell data types

In the second part of developing SMILE, we demonstrate that SMILE can handle most types of single-cell omics data. Besides RNA-seq data shown above, we tested SMILE in ATAC-seq data from Mouse ATAC Atlas and a Hi-C data from mouse embryo cells (Cusanovich et al. 2018; Collombet et al. 2020). SMILE can distinguish tissue types within Mouse ATAC Atlas, and it also recovers most cell types in the brain tissue (Supplementary Fig. S2A). For clustering single cell Hi-C data, SMILE has a slight advantage of distinguishing different cell stages compared to PCA (the baseline) (Supplementary Fig. S2B). However, we want to point out that, for a single source data, SMILE does not show substantial difference from a standard PCA approach. Since PCA finds most variations and it is unlikely that unwanted variations are confounded within a single-source data, there would be no obvious advantage of using SMILE to find biological variations, and we would not recommend users to use SMILE for single-source data.

### SMILE eliminates batch effects in data from multiple sources

It is now common to find multiple single-cell transcriptomics datasets for the same tissue or biological system generated by different techniques or research groups. A standard clustering analysis often fails to identify cell types, but instead only detects differences between experimental batches. In contrast, SMILE directly learns a representation that is not confounded by batch effect and can be combined with common clustering methods for cell type identification. We tested batch-effect correction in human pancreas data, human peripheral blood mononuclear cell (PBMC) data, and human heart data (Baron et al. 2016; Grün et al. 2016; Muraro et al. 2016; Segerstolpe et al. 2016; Lawlor et al. 2017; Zheng et al. 2017; Litviňuková et al. 2020; Tucker et al. 2020). In this study, we compared SMILE with Harmony, which has been reported as a top method for batch-effect correction (Korsunsky et al. 2019; Tran et al. 2020). We showed that SMILE has comparable performance to Harmony in these 3 systems (Fig. 1B-D) and both removed batch effects and recovered cell types identified in original reports (Fig. 1E). Meanwhile, the integrated representations learned by SMILE and Harmony are classification-friendly for label transferring (Fig. 1F).

### Data integration across multiple tissues

One advantage of SMILE in contrast to Harmony is that SMILE can perform batch-effect correction without needing explicit batch information to be provided. This is useful when batch information is complicated by other differences, as is the case for data collected from different tissues: these experiments are different both biologically and in their technical batch effects. We next tested SMILE on such a dataset: single-cell transcriptome data from 6 different tissues in Human Cell Landscape (Han et al. 2020). Without dealing with batch effects and sample differences, a standard analysis pipeline recovers tissue differences as the major source of variation, while endothelial cells, which we focus on as an example cell type, are spread out rather than clustered within each tissue (Fig. 2A). However, by applying SMILE to these tissues, we show that SMILE can find a common representation for cell types (e. g. endothelial cell, B cell, and T cell) that are conserved across these tissues (Fig. 2B). This is similar to an ability recently reported for the MARS approach, but, in contrast to MARS, SMILE is trained in a fully unsupervised manner (Brbić et al. 2020).

**Fig. 2.**
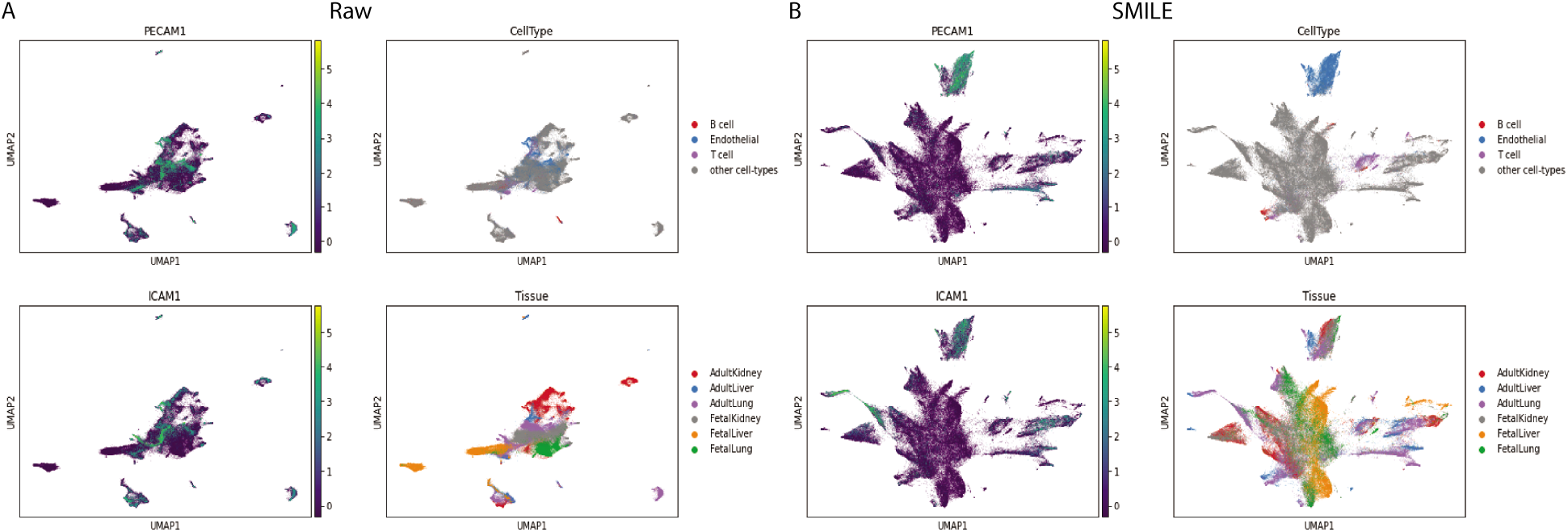
Integration of 6 tissues of Human Cell Landscape transcriptomic data. (A) Visualization without integration: inputting raw data into UMAP. Expression of two endothelial markers (PECAM1 and ICAM1) are visualized in the 2D UMAP (left column). Cells are colored either by cell-types or tissue-types and visualized in the 2D UMAP (right column). (B) SMILE integration. Expression of two endothelial markers (PECAM1 and ICAM1) are visualized in the integrated 2D UMAP (left column). Cells are colored either by cell-type or tissuetype and visualized in the integrated 2D UMAP (right column).

### Architecture of p/mpSMILE for integrating joint profiling omics data

To apply SMILE to joint profiling data, we modified it into new architectures, pSMILE (Supplementary Fig. S3A) and mpSMILE (Fig. 3A). These new architectures contain two separate encoders (Encoder A and Encoder B). Encoder A projects RNA-seq or methylation data into representation *z^a^*, while Encoder B handles projection of ATAC-seq or Hi-C into representation *z^b^*. We aim to learn *z^a^* and *z^b^* that will be confined in the same latent space, so we apply the same one-layer MLPs to each, which further reduce them into 32-dimension output and probabilities of *K* pseudo cell types. Using the RNA/ATAC- or Methyl/Hi-C-joint data to train SMILE would be the same as using the self-paired data, except that we introduced two separate encoders. We tested the performance of pSMILE on a simulated joint single-cell transcriptome dataset and two joint profiling datasets generated by SNARE-seq and sci-CAR (Supplementary Fig. S3B-C and Supplementary Fig. S4) (Cao et al. 2018; Chen et al. 2019). The results with simulated joint data, produced by splitting a single scRNA-seq dataset into two subsets with separate genes indicates that pSMILE can integrate data from two entirely different feature spaces (Supplementary Fig. S3C). Our pSMILE results with joint RNA-seq and ATAC-seq data indicate that RNA-seq data have a greater cell type discriminative power than ATAC-seq, so we created the further modified mpSMILE with two duplicated Encoder As that share the same weights and are used to amplify the effect of RNA-seq datasets. With the discriminative power of RNA-seq, mpSMILE revealed more cell types in the mixed cell line and mouse kidney data (Supplementary Fig. S5). As in these examples, mpSMILE successfully integrates the joint profiled data even with mismatched feature spaces: RNA-seq is quantified per gene and ATAC-seq is represented by genome wide peaks that are not linked to specific genes.

**Fig. 3.**
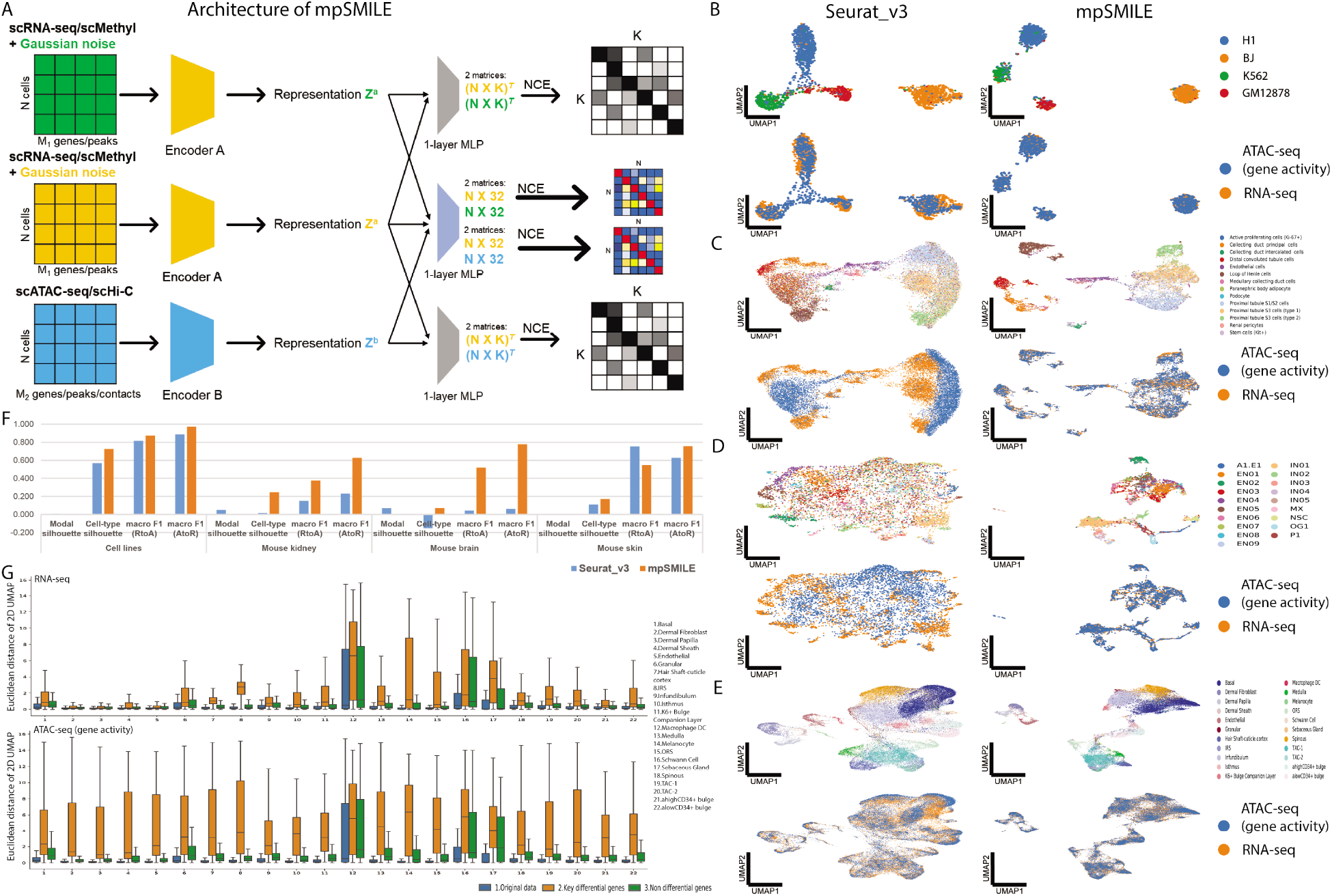
Integration of scRNA-seq and scATAC-seq through p/mpSMILE. (A) Architecture of mpSMILE. scRNA-seq or scMethyl data are forwarded through Encoder A to produce representation *z^a^*, and scATAC-seq or scHi-C are forwarded through Encoder B to produce representation *z^b^*. mpSMILE has duplicated Encoder As, and cells in scRNA-seq/scMethyl part would be duplicated by adding gaussian noises and become self-pairs, as we did in SMILE for multi-source single cell transcriptome data. Two one-layer MLPs in mpSMILE are the same as those in SMILE. (B-E) UMAP visualization of integrated representation of (B) mixed cell-lines, (C) mouse kidney, (D) mouse brain and (E) mouse skin data, by Seurat_v3 (left) and mpSMILE (right). Cells are colored by cell-type in the upper panels and colored by data types in the lower panels. (E) Evaluation of integration by macro F1, modal, and cell-type silhouette scores. For macro F1 and cell-type silhouette score, 1 indicates the best performance, and higher is better. RtoA represents label transferring from RNA-seq to ATAC-seq, and AtoR represents from ATAC-seq to RNA-seq. For modal silhouette score, 0 is the best. (F) boxplot of Euclidean distances between paired cells. Blue box: original data was forwarded through trained mpSMILE and Euclidean distances between cells in RNA-seq and their corresponding cells in ATAC-seq were measured in the integrated 2D UMAP. Orange box or green box: either key differential genes or non-key genes were suppressed to zeros, then the suppressed data was forwarded through trained mpSMILE, and Euclidean distances between cells in RNA-seq and their corresponding cells in ATAC-seq were measured in the integrated 2D UMAP. Upper panel is suppression of key gene expression, and lower panel is suppression of key gene activity.

### Joint clustering through mpSMILE improves upon previous methods and reveals key biological variables

In projecting joint profiling data into the same space, SMILE accomplishes a similar purpose as Seurat_v3, but with more flexibility of input. Seurat_v3 implements canonical correlation analysis (CCA) to project both RNA-seq and ATAC-seq data into the same low-dimension space (Stuart et al. 2019). The use of CCA requires the two datasets to share the same features. As shown above, SMILE can work with datasets where the two data types involve entirely different features (e.g. genes vs. genomic bins). To compare SMILE with Seurat_v3, we re-quantified the ATAC-seq into gene activities, and we further included mouse brain and mouse skin datasets generated by SHARE-seq (Ma et al. 2020). Since cell pairs are known in these datasets, we used all pairs to train both Seurat_v3 and mpSMILE. We visualized integration results of Seurat_v3 and mpSMILE by UMAP. Both Seurat_v3 and mpSMILE were able to project ATAC-seq and RNA-seq data into the same space while discriminating between cell types for the mixed cell-lines data (Fig. 3B). However, Seurat_v3 showed poor performance on the mouse kidney data by sci-CAR and the mouse brain data by SHARE-seq, failing to project the two data sources into the same space and failing to distinguish cell types (Fig. 3C and D). For the mouse skin data by SHARE-seq, Seurat_v3 and mpSMILE have comparable performance (Fig. 3E). Compared to Seurat_v3 in a quantitative way, SMILE shows higher F1 scores for either transferring RNA-seq label to ATAC-seq or ATAC-seq to RNA-seq and better multimodal integration in terms of modal and celltype silhouette scores (Fig. 3F).

To evaluate which biological factors drive the co-embedding we observe, we set candidate genes from the mouse skin to zero and passed this altered data through the mpSMILE encoder to evaluate whether co-embedding would be disrupted. Indeed, when we remove key differential genes (Supplementary Fig. S6), clusters are greatly disrupted in the co-embedding (Fig. 3G and Supplementary Fig. S7). Since this evaluation does not require re-training, this approach would allow rapid screening to detect which gene expression and chromatin accessibility features best explain the cell-type separation.

Though p/mpSMILE was designed to do joint clustering for joint profiling data, it can be combined with pair-identification tools to achieve integration for non-joint-profiling data. Seurat_v3 implements “FindTransferAnchors” function, which can identify quality pairs in bimodal datasets. Here, we combined Seurat_v3 and SMILE to achieve integration for non-joint-profiling PBMC data. Empowered by Seurat_v3, SMILE did decent integration for PBMC data (Supplementary Fig. S8). Notably, after pairing is accomplished, SMILE allows the user to input mismatched features for the two modalities. Here, for example, ATAC-seq data was used at the peak level while RNA-seq data was used at the gene level. Thus, differences between cell type discrimination by SMILE as compared to Seurat may represent real differences in the information captured by genomewide peaks instead of only activities assigned to genes.

### Application of p/mpSMILE in joint profiling DNA methylation and chromosome structure data

We next evaluated the applicability of SMILE to the integration of joint profiling DNA methylation and chromosome structure data of mESC and NMuMG cells, and the human prefrontal cortex (PFC). Unlike integration of RNA-seq and ATAC-seq, it is difficult to match DNA methylation features to chromatin structure features. Therefore, using CCA for integration of Methyl and Hi-C as in Seurat_v3 would be a challenging task. In such a case, SMILE has the unique advantage of not requiring feature matching. We applied pSMILE in both mESC and NMuMG data and human PFC data. pSMILE can distinguish mESC and NMuMG cells but only revealed 5 major cell types in human PFC (Fig. 4A and B). Then, we applied mpSMILE by using Methyl data in place of RNA-seq and Hi-C in place of ATAC-seq, because Methyl data recovers more distinct cell types in (Lee et al. 2019). However, mpSMILE did not reveal more cell types than pSMILE (Fig. 4B). Because we used all 100kb bins of CG methylation as input for SMILE, it is possible that SMILE cannot fully unlock discriminative power of methylation data. Thus, we further projected Hi-C cells onto the tSNE space of CG methylation from the original study, but in a SMILE manner. The tSNE space of CG methylation preserves the distinct structure for each cell type identified by the author, and it should save us from learning discriminative representation for Methyl data. However, neuron sub-types did not align well with their methylation counterparts in this projection task, though other cell types did (Fig. 4C). These results suggest that the Hi-C part of human PFC data lacks discriminative power to distinguish neuron sub-types as the original report showed. This is also an important demonstration that SMILE will not force datasets to overlap if they are truly capturing different biological features and there is a gap between the two modalities.

**Fig. 4.**
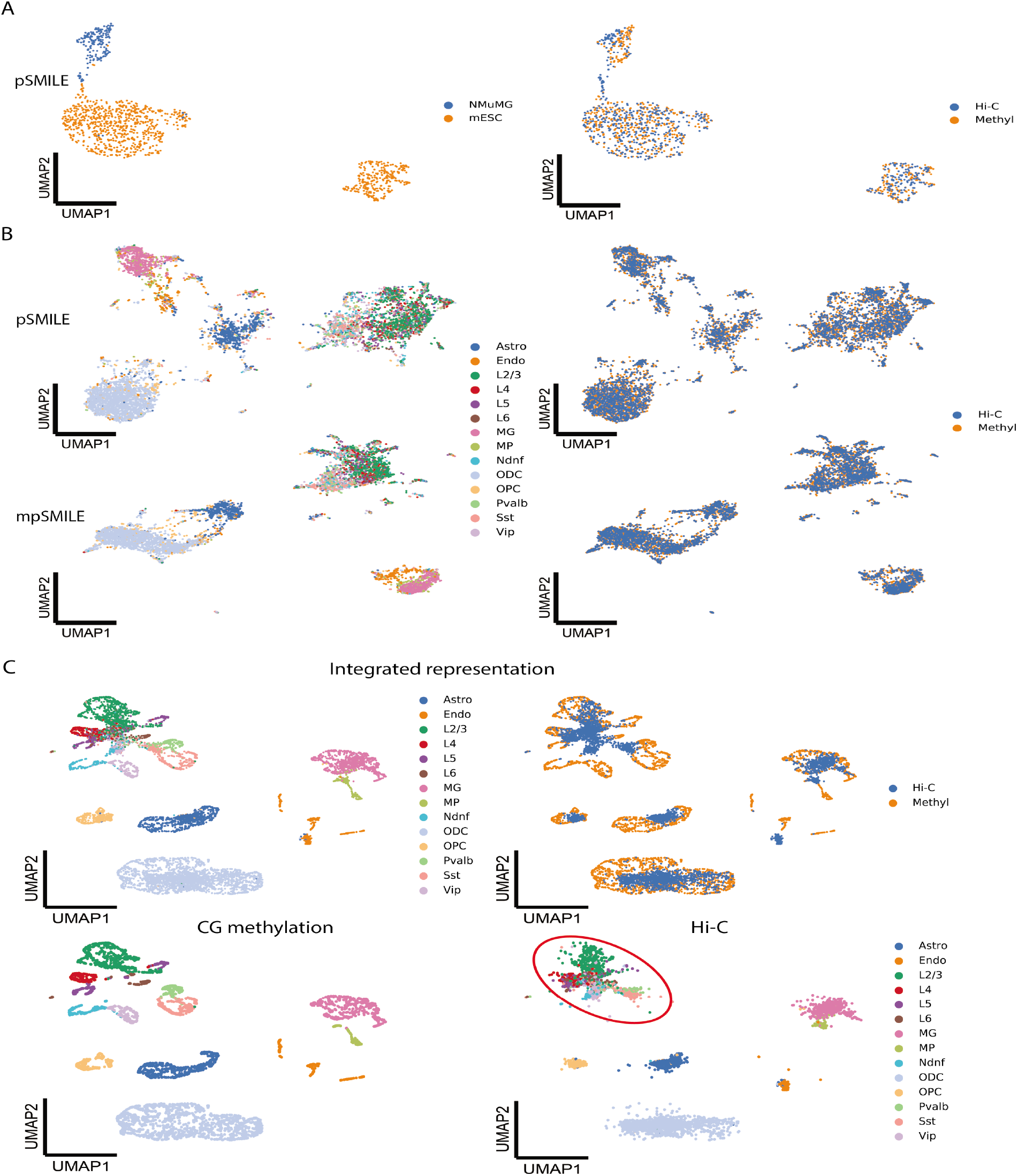
Integration of scMethyl and scHi-C through p/mpSMILE. (A and B) UMAP visualization of integrated representation of A) mESC and NMuMG cells and B) human PFC data, by p/mpSMILE. Cells are colored by cell-types reported by the author (left panel), or data types (right panel). (C) Projection of Hi-C onto tSNE space of CG methylation using SMILE. We used the tSNE space of CG methylation as input for Encoder A instead of 100kb bins of CG methylation. Training SMILE in this case becomes training Encoder B to project Hi-C data into the tSNE space of CG methylation, though we visualized the integrated representation through UMAP. Top row: Hi-C and methylation data on the same graph. Second row: Same representation as above, but with CG methylation (left) and Hi-C (right) shown separately. Red circle highlights region of indistinct neuronal cell types.

### Combining SMILE and pSMILE for integration of more than 2 data modalities

The recently published Paired-Tag technology can jointly profile one histone mark and gene expression from the same nucleus (Zhu et al. 2021). The unique design of this study paired RNA-seq data with 5 different histone marks, and it provides us demonstration data to show how we can combine SMILE and pSMILE to achieve integration of more than 2 modalities. With these modifications, SMILE can integrate RNA-seq, H3K4me1, H3K9me3, H3K27me3, and H3K27ac (Fig. 5A). In the first step, we used SMILE to integrate RNA-seq data from 6 batches, as we did previously for multi-source transcriptome data. Then, we replaced Encoder A in pSMILE with the trained encoder in SMILE with frozen weights. Therefore, RNA-seq data would be only forwarded through the Encoder A to generate representation *z^a^* and no gradients will be sent back during training. Since SMILE had already learned discriminative representation for RNA-seq data, training pSMILE in the second step was aimed to project histone mark data into the representation of RNA-seq. Because these histone mark data are not paired, we trained 4 pSMILE models with 4 paired RNA-seq/Histone marker data. Finally, we can project all nuclei from the 5 modalities into the same UMAP space for visualization. Indeed, this approach preserved distinct properties of cell types while mixing data types (Fig. 5B). As we did above with the ATAC-seq and RNA-seq joint profile, this learned encoder could be used to screen for which modification peaks are most important for cell type discrimination.

**Fig. 5.**
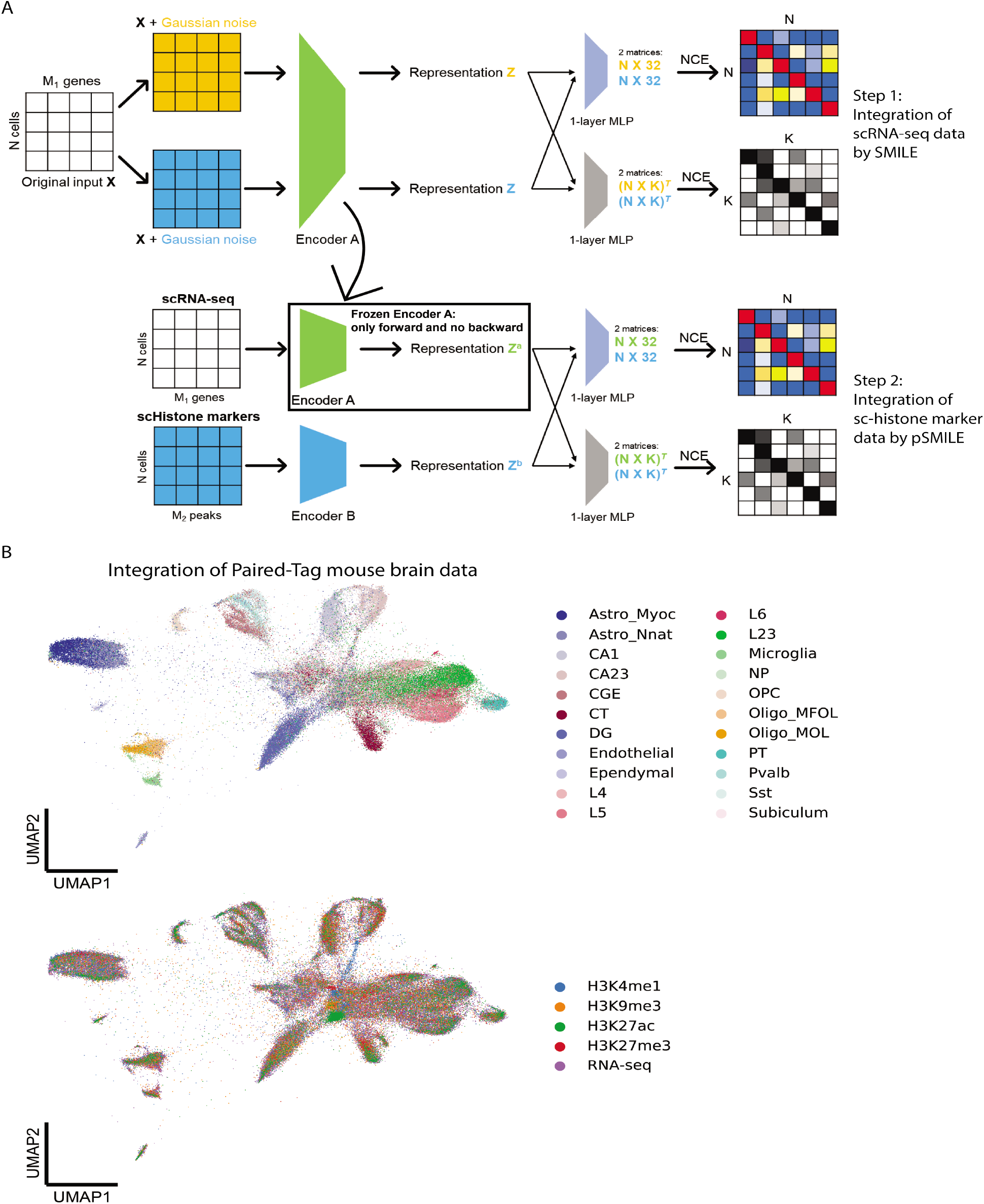
Integration of Paired-Tag mouse brain data through SMILE. (A) Procedure of combining SMILE and pSMILE for integration of Paired-Tag data. (B) UMAP visualization of integrated representation of mouse brain data. Cells are colored by cell-types (upper panel) and data types (lower panel).

## Discussion

In this study, we presented three variants of SMILE models that perform single cell omicd data integration tasks in different cases, and our SMILE approach effectively accomplished these tasks with comparable or even better outcomes than existing tools. In multi-source single-cell transcriptome data, we demonstrated that SMILE removes batch effects as well as Harmony. In multimodal single-cell data, SMILE saves users from extra feature engineering while performing integration. Encoders learned by SMILE can be used to determine what biological factors underlie the derived joint clustering and to transfer cell type labels to future related experiments. Finally, we demonstrated how to combine or modify our SMILE models to address the integration of more than 2 modalities. For the joint-profiling data and the Paired-Tag data, the learned encoders could be used to project other single source datasets (e.g. ATAC-seq, ChIP-seq, or Hi-C without paired RNA-seq or Methylation data) into this same space to classify cell types and compare cell type distributions in different conditions. In the case of the representation we learned from Paired-Tag for example, this would mean that a future experiment that measures only H3K27Ac in single cells could be projected into this comprehensive histone mark and transcriptome-defined representation space for cell type classification and comparison.

Overall, our SMILE approach shows the ability to integrate single-cell omics data as a comprehensive tool. One limitation of SMILE for multimodal integration is that cell pairs must be known. Therefore, training SMILE involves creating self pairs across a single modality or using the natural pairs in joint profiling data. But we combined Seurat_v3 and SMILE to show how SMILE can also be used for non-joint profiling data. Future work in SMILE development can incorporate strengths of the other two new data integration approaches, scJoint (Lin et al. 2021) and GLUER (Peng et al. 2021) to enable SMILE to better handle integration of non-joint profiling data. As joint profiling technologies become a new mainstream in single-cell research, we envision that SMILE will continue to reveal its potential in data integration.

## Methods

### Architecture of SMILE, p(paired)SMILE and mp(modified paired)SMILE

All three variants of SMILE have encoders as the main components for feature extraction. In SMILE, there is only one encoder. This encoder consists of two fully connected layers that have 1000 and 128 nodes, respectively. Each fully connected layer is coupled with a BatchNorm layer to normalize output which is further activated by ReLu function. Different from SMILE, pSMILE and mpSMILE have another encoder that has the same structure as the one in SMILE but takes an input with different dimension. The use of two independent encoders (Encoder A and B) in pSMILE and mpSMILE is to handle inputs from two different data sources with different features, for example RNA-seq and ATAC-seq. Therefore, pSMILE and mpSMILE do not require extra feature engineering to match features for inputs from two different data sources. Modified from pSMILE, mpSMILE has a duplicated Encoder A, which shares the same weights. Using duplicated encoders takes advantage of the discriminative power of RNA-seq or Methyl to learn a more discriminative representation. This is because we observed a compromise between the source with more discriminative power and the source with less. We stack two independent fully connected layers to the encoder(s), which further reduce the representation *z* to a 32-dimension vector with ReLu activation and *K* pseudo cell-type probabilities with SoftMax activation. In our application of SMILE, we set *K* to 25, or in the case of semi-supervised learning, *K* can be set to the number of known cell types.

### Cell pairing

SMILE takes paired cells as inputs. When using SMILE for integration of multi-source single-cell transcriptome data, we treat each cell itself as positive pair. To prevent the two cells in each pair from being completely the same, we add gaussian noise to differentiate them. Thus, the learning process will maximize the true similarities between the pair while minimizing the effect of noise. When using pSMILE and mpSMILE for integration of multimodal single-cell data, we pair a cell from RNA-seq/Methyl with its counterpart from ATAC-seq/Hi-C. Since joint profiling quantifies two aspects of one single cell, we know that one cell in RNA-seq/Methyl has a corresponding cell in ATAC-seq/Hi-C. When RNA-seq and ATAC-seq come from two separate studies, the user may need to pair cells of the same cell type manually. Here, we suggest using “FindTransferAnchors” function in Seurat_v3 to generate cell pairs. Then, user can use these paired cells to train p/mpSMILE.

### Loss function

a. **Noise-contrastive estimation (NCE).** The core concept of making cells resemble themselves resides in the use of NCE as the main loss function. In training, we divide a whole dataset into multiple batches, and each batch has *N* cells. For multi-source singlecell transcriptome data, we differentiate each cell into two by adding random gaussian noise. Therefore, there are 2*N* cells in one batch. In each batch, each cell has itself as positive sample and the rest of 2(*N* – 1) cells as its negative samples. For joint profiling data and in an *N*-pair batch, one of *N* cells in RNA-seq/Methyl has its corresponding cell among *N* in ATAC-seq/Hi-C as the positive sample and the rest of 2(*N* – 1) cells summed from both RNA-seq/Methyl and ATAC-seq/Hi-C as negative samples. Let *sim* (*u, v*) = denote the dot product between *L*_2_ normalized *u* and *v*. Then, NCE for a positive pair of examples (*i,j*) can be defined as (Eq. 1). In this study, we applied NCE to 32-dimension outputs (two matrices in a shape of *N* by 32) with *τ* as 0.05 and *K* pseudo cell-type probabilities (two matrices in a shape of *K* by *N*) with *τ* as 0.15.
b. **Mean squared error (MSE).** We use MSE as additional loss function in p/mpSMILE to push the representation of ATAC-seq/Hi-C to be closer to the representation of RNA-seq/Methyl in the latent space (Eq. 2). Of note, MSE alone is unable to drive the model to learn a discriminative representation (Supplementary Fig. S9 and Fig. S10).

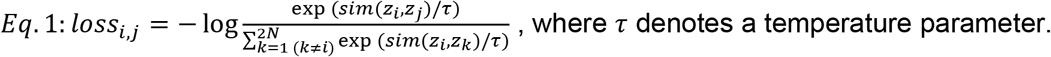

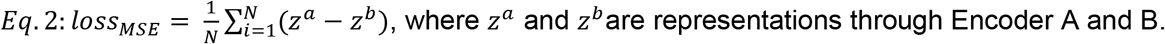

### Evaluation of data integration

To evaluate batch-effect correction, we use ARI as the evaluation metric. We perform Leiden clustering to re-cluster cells using the batch-removed representation, and we compare the new clustering label with the cell-type label reported by the authors. For fair comparison of SMILE and Harmony, we use multiple resolutions to get the clustering label that has the highest ARI against authors’ cell-type label, and report that ARI value as the performance of SMILE or Harmony in that data.

To evaluate integration of multimodal single-cell data, we select silhouette score as the metric. We defined two silhouette scores based on two sets of labels. For modal silhouette, the score was calculated using data type label. Closer to 0, better mixing of RNA-seq and ATAC-seq. For cell-type silhouette, the score was calculated using cell-type label. Closer to 1, better preservation of distinction of different cell-types. Meanwhile, we know each cell pair because of joint profiling. So, we also measure Euclidean distance of paired cells in the 2D UMAP space, before and after training, to show if paired cells become closer in the integrated representation.

### Evaluation of label transferring

Once we have an integrated representation for multiple datasets, we can transfer labels from known data to other unknown data. To evaluate label transferring, we use weighted F1 and macro F1 scores, implemented in sklearn. In integration of multi-source single-cell transcriptome data, we select one source as the training set to train a Support Vector Machine (SVM) classifier, and test the accuracy measured through weighted and macro F1 scores in other sources. In joint profiling data, we can use the cell-type label as ground-truth to train a SVM classifier on RNA-seq or Methyl and test it in ATAC-seq or Hi-C. Vice versus.

### Processing of RNA-seq, ATAC-seq, Methyl, Hi-C, and histone marker data

For RNA-seq data, raw gene expression count data was normalized through a ‘LogNormalize’ method, which normalizes the raw count for each cell by its total count, then multiplies this by a scale factor (10,000 in our analysis), and log-transforms the result. Then, we use “highly_variable_genes” function in Scanpy to find most variable genes as input for SMILE (Wolf et al. 2018). For ATAC-seq data at peak level, we perform TF-IDF transformation and select top 90-percentile peaks as the input for SMILE. For ATAC-seq data at gene level, we first use “CreateGeneActivityMatrix” function in Seurat_v3 to sum up all peaks that fall within a gene body and its 2,000bp upstream, and we use this new quantification to represent gene activity (Stuart et al. 2019). Then, we apply LogNormalize to gene activity matrix and find most variable genes as input for SMILE. For CG methylation data, we first calculate CG methylation level for all nonoverlapping autosomal 100 kb bins across entire human genome. Then, we apply LogNormalize to the binned CG methylation data. For Hi-C data, we use scHiCluster with default setting to generate an imputed Hi-C matrix at 1MB resolution for each cell (Zhou et al. 2019). Due to the size of Hi-C matrix, we are unable to concatenate all chromosomes to get a genome-wide Hi-C matrix. Therefore, we use a dimension-reduced Hi-C data of whole genome, which is implemented in scHiCluster. For histone marker data, we perform TF-IDF to transform the raw peak data and select top 95-percentile peaks as the input for SMILE.

### Data integration through SMILE/pSMILE/mpSMILE

We used “StandardScaler” in sklearn to scale all input data before training SMILE, except that the dimension-reduced Hi-C data has been scaled. We trained SMILE with batch size as 512 for all multi-source single-cell transcriptome data in this study, and the SMILE model can converge within 5 epochs in all cases, indicated by the total loss. In integration of 4 joint profiling RNA-seq and ATAC-seq data, we use all cell pairs for training, and we trained p/mpSMILE for 20 epochs with batch size as 512. For sn-m3C-seq data, we trained p/mpSMILE on whole data for 10 epochs with batch size as 512. For all experiments in this study, we used learning rate as 0.01 with 0.0005 weight decay.

### Identification of key and non-key differential genes

We performed wilcoxon test to identify key differential genes and their ranking in both mouse skin RNA-seq and ATAC-seq data, using cell-type label from the original report. In both tests, we identified most genes are significantly differentially expressed (RNA-seq) or active (ATAC-seq), since these genes are also highly variable. Thus, we only selected top 15 genes in RNA-seq and top 150 genes in ATAC-seq of each cluster as key differential genes. This resulted 238 key differential genes against 361 non-key genes in RNA-seq and 1369 key genes against 1914 non-key genes in ATAC-seq. For testing which features contribute to the representation learned by SMILE, key genes for each cluster were sequentially assigned values of zero and then the dataset was fed back through the encoder to determine the effect on the representation. This is conceptually similar to the motif screening used to probe a deep learning representation in (Fudenberg et al. 2020).

### Batch removal through Harmony

We use Harmony with default setting to integrate multi-source single-cell transcriptome data, and the code can be found at https://pypi.org/project/harmony-pytorch/. Though we did not observe much difference between SMILE and Harmony, in terms of batch-effect correction, we noticed that SMILE ran much faster than Harmony, because of use of GPU computation in training SMILE (Supplementary Table S1).

## Data integration through Seurat_v3

We follow a default workflow of Seurat_v3 to do the integration of SHARE-seq mouse skin data. The workflow code can be found at https://satijalab.org/seurat/v3.2/atacseq_integration_vignette.html. For a fair comparison, we did not use “FindTransferAnchors” function to find anchors. Instead, we used all pairs as we did for SMILE. Then, we used these paired cells as anchors for Seurat to learn the co-embedding of RNA-seq and ATAC-seq.

**Supplementary Table S1.** Running time of SMILE and Harmony on human pancreas, human PBMC, human heart and 6 tissues of Human Cell Landscape data.

| Dataset | Number of cells | Runtime (min) |  |
| --- | --- | --- | --- |
|  |  | SMILE | Harmony |
| Human pancreas | 14,890 | 1 | 2.5 |
| Human PBMC | 15,476 | 1 | 2.5 |
| Human heart | 646,921 | 8 | 30 |
| Human Cell Landscape (6 tissues) | 107,829 | 3 | NA |

## Data availability

All data used in this study are publicly available with GEO accession (Supplementary Table S2). Some studies have processed data that are hosted by institution data portals (Supplementary Table S2).

**Supplementary Table S2.**
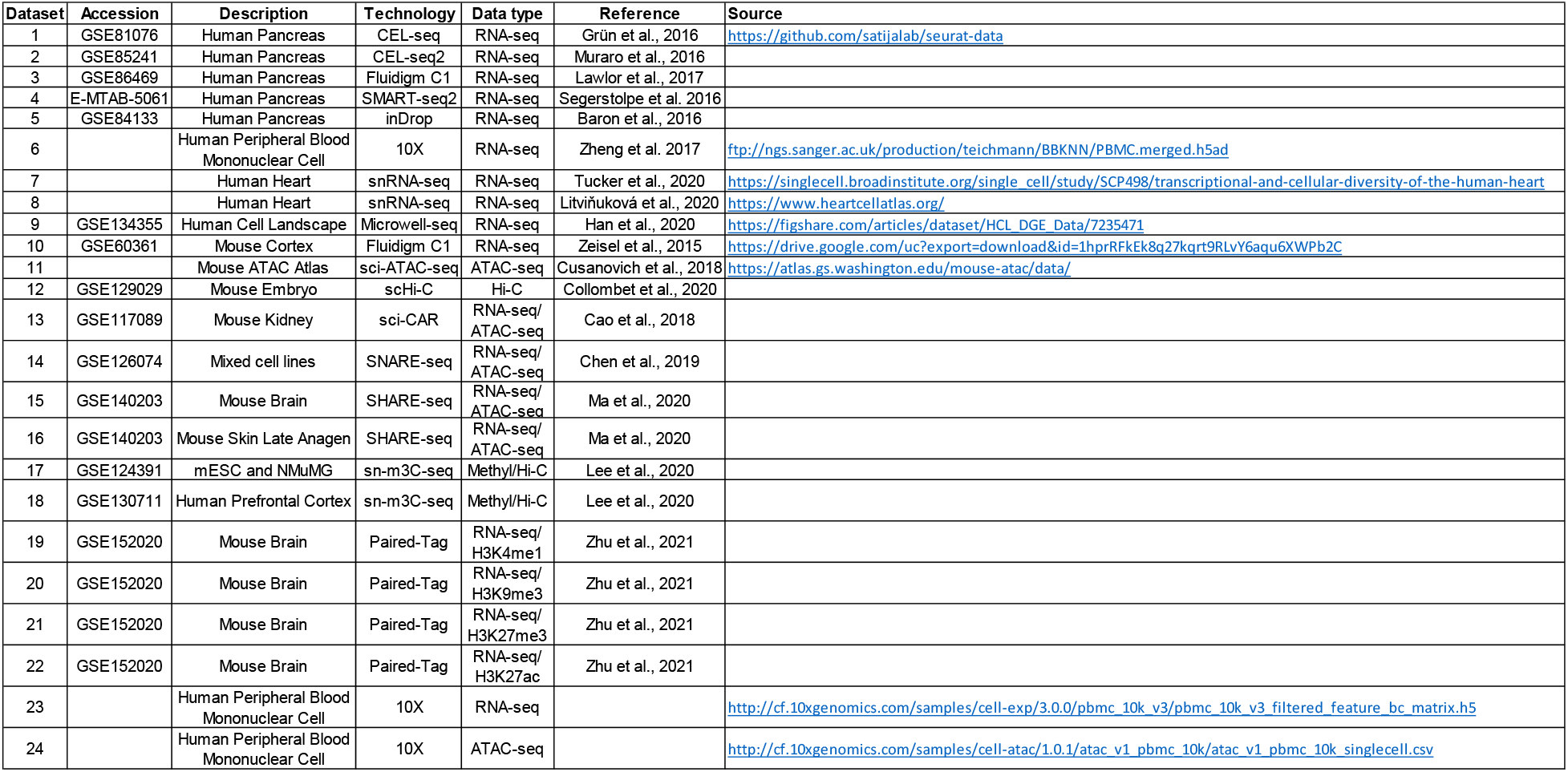
Summary of data sources used in this study, including GEO accession, brief description, and link to the processed data if applicable.

## Code availability

The source code of SMILE including analyses of key results in the study can be found at: https://github.com/rpmccordlab/SMILE.

## Author Contributions

Y.X. conceived and developed the method with guidance from R.P.M. and produced all the figures. P.D. computationally processed raw sequencing data for input to SMILE. Y.X. and R.P.M. wrote the manuscript with input from P.D.

## Acknowledgements

We thank Joseph Ecker’s lab for sharing information about processing their sn-m3C-seq data. This research was supported in part by NIH NIGMS grant R35GM133557 to R.P.M.

