## Supplementary Figures for "SMILE: Mutual Information Learning for Integration of Single Cell Omics Data"

Figure S1

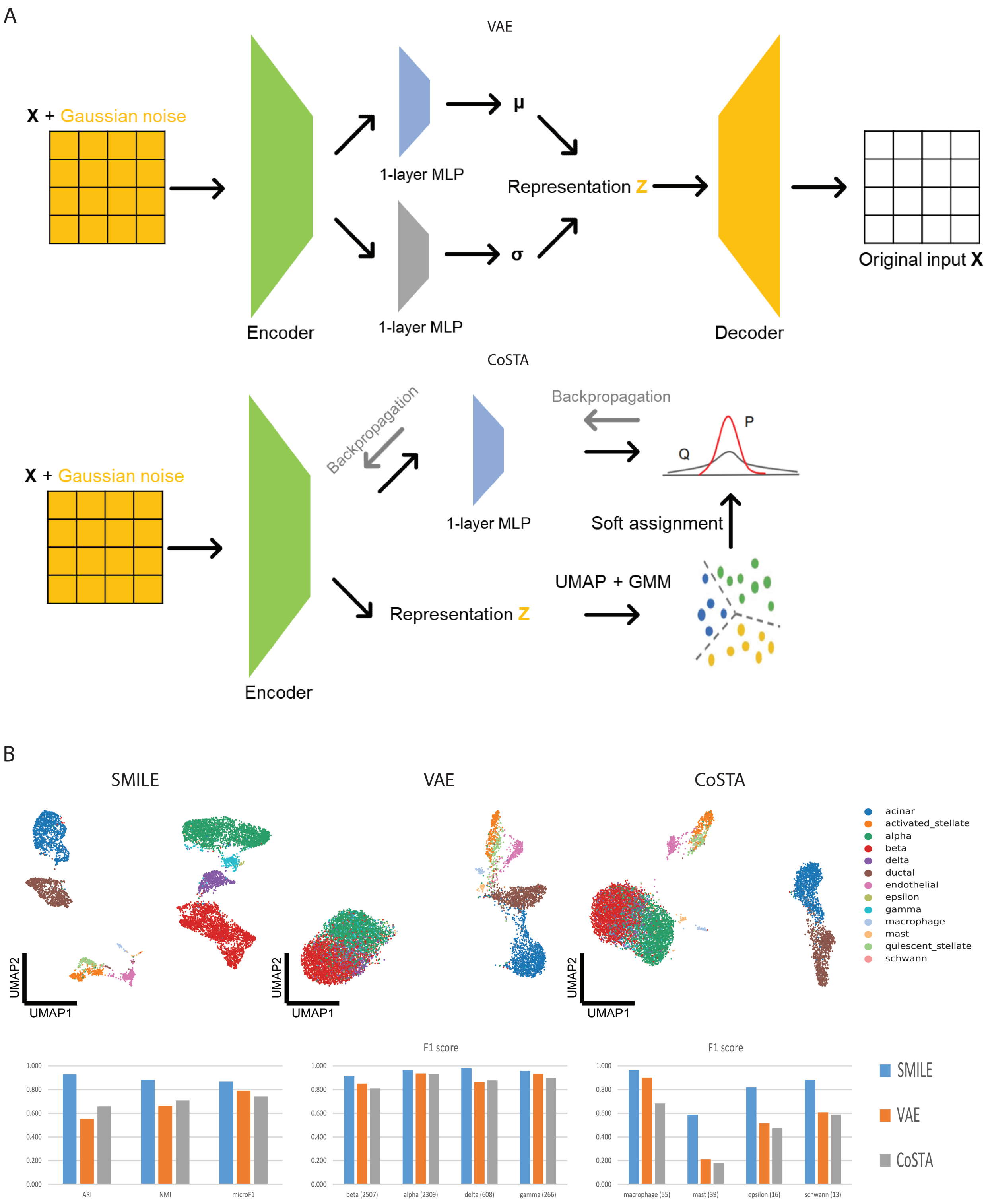


**Supplementary Fig. S1**

Comparison of SMILE, VAE and CoSTA. a, Architectures of VAE and CoSTA. Encoders in VAE and CoSTA have the same structure as the encoder in SMILE. b, UMAP visualization of representation of a single-source human pancreas data and numerical comparison in terms of recovering major and minor cell-types (F1 scores).

Figure S2

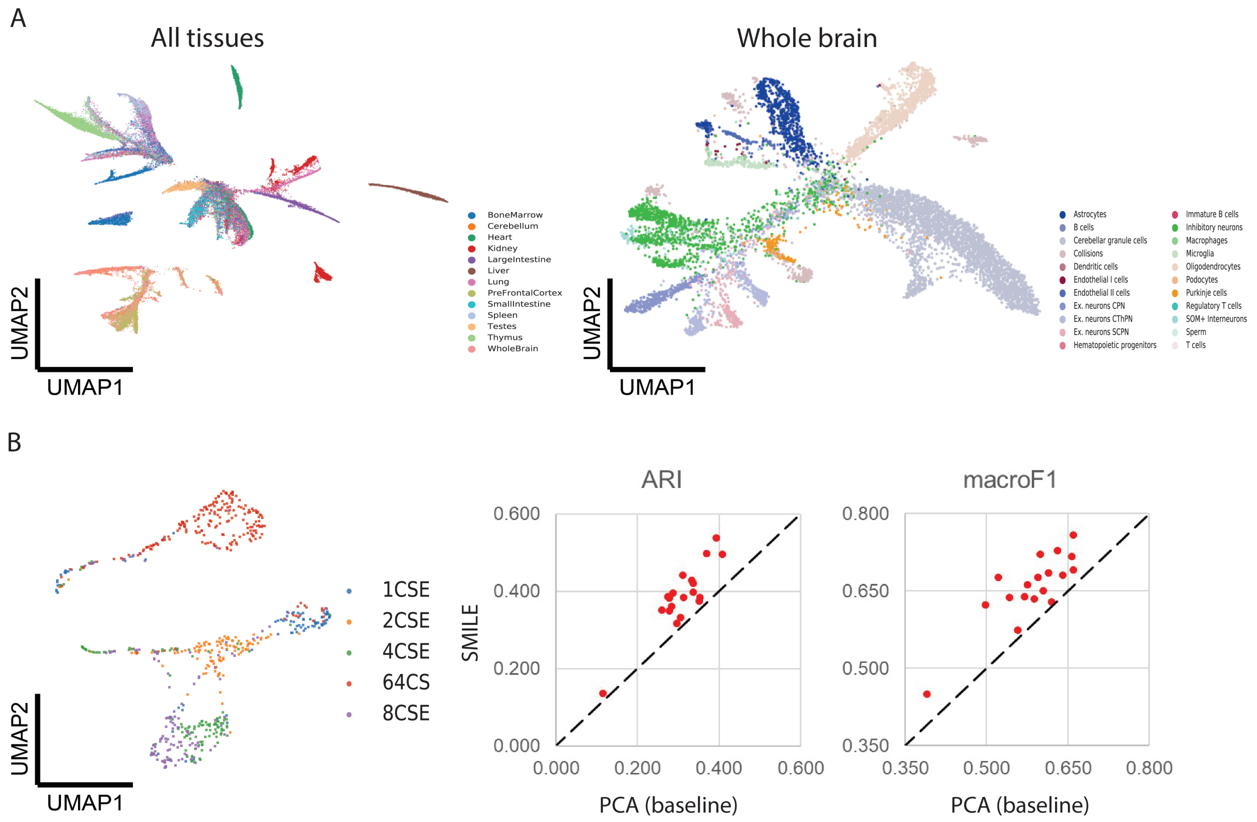


**Supplementary Fig. S2**

Application of SMILE in single-cell ATAC-seq and Hi-C. a, UMAP visualization of SMILE representation of Mouse ATAC Atlas. Left panel is the visualization of whole dataset and cells are colored by tissue types. Right panel is the visualization of a subset of cells from mouse brain. Cells are colored by cell-types reported by the author. b, UMAP visualization of SMILE representation of mouse embryo single cell Hi-C data (left), and comparison of SMILE and PCA (baseline) for each different chromosome separately (right). Cells are colored by developmental stages. Calculation of ARI and macro F1 is based on the developmental stage of a cell as the ground truth.

Figure S3

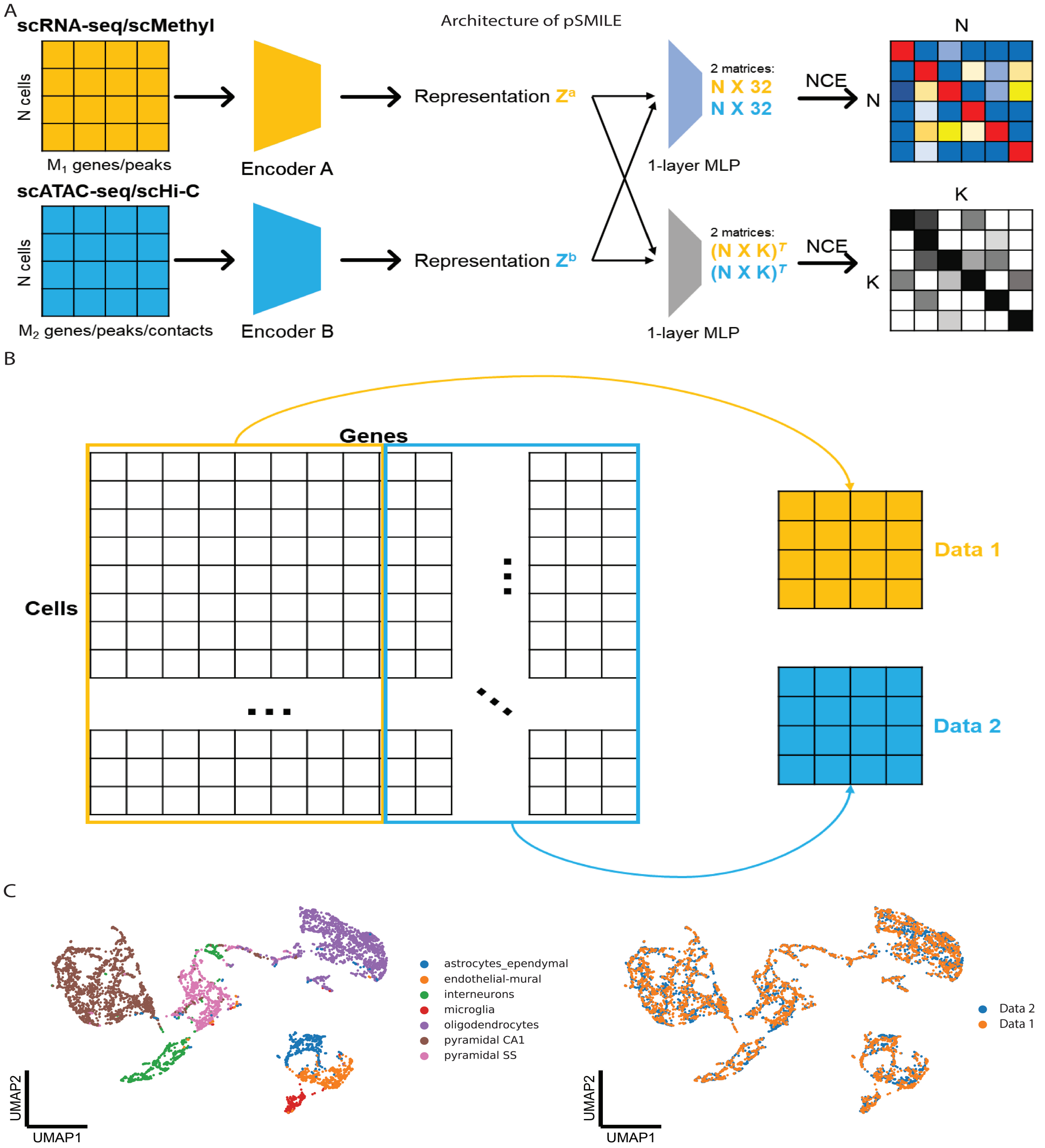


**Supplementary Fig. S3**

Integration of synthetic multimodal single-cell omics data through pSMILE. a, Architecture of pSMILE. scRNA-seq/scMethyl would be forwarded through Encoder A to produce representation $z^a$, and scATAC-seq/scHi-C would be forwarded through Encoder B to produce representation $z^b$. Two one-layer MLPs in pSMILE are the same as those in SMILE. b, Construction of synthetic multimodal single-cell data. The synthetic multimodal data is based on a real single-cell RNA-seq data from mouse cortex.(Zeisel et al. 2015) Data 1 and data 2 have the same cells, and each cell in data 1 is paired with its corresponding cell in data 2. Data 1 and data 2 are generated from the original data through splitting genes into two halves. Therefore, data1 and data 2 do not share any common features. c, UMAP visualization of integrated representation of mouse cortex by pSMILE. Cells are colored by cell-types (left) and data types (right).

Figure S4

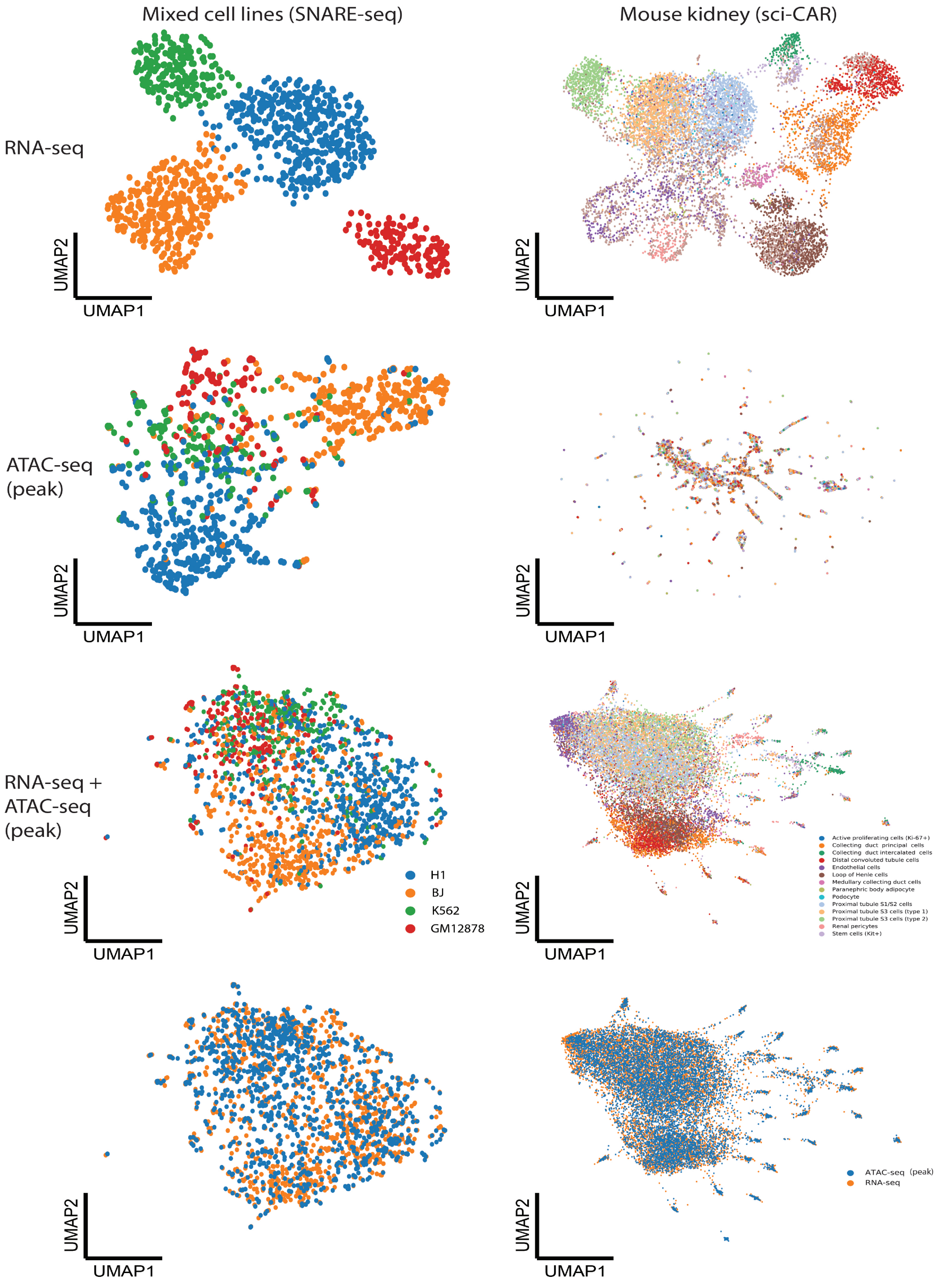


**Supplementary Fig. S4**

Integration of SNARE-seq (left) and sci-CAR (right) data through pSMILE. The first two rows show independent visualization of RNA-seq and ATAC-seq at peak level. Cells are colored by cell-types reported in original studies. The 3$^{rd}$ and 4$^{th}$ rows show the UMAP visualization of integrated representation of mixed cell line data (left) and mouse kidney data (right) by pSMILE. Cells are colored by cell-types (3$^{rd}$ row) and data types (4$^{th}$ row).

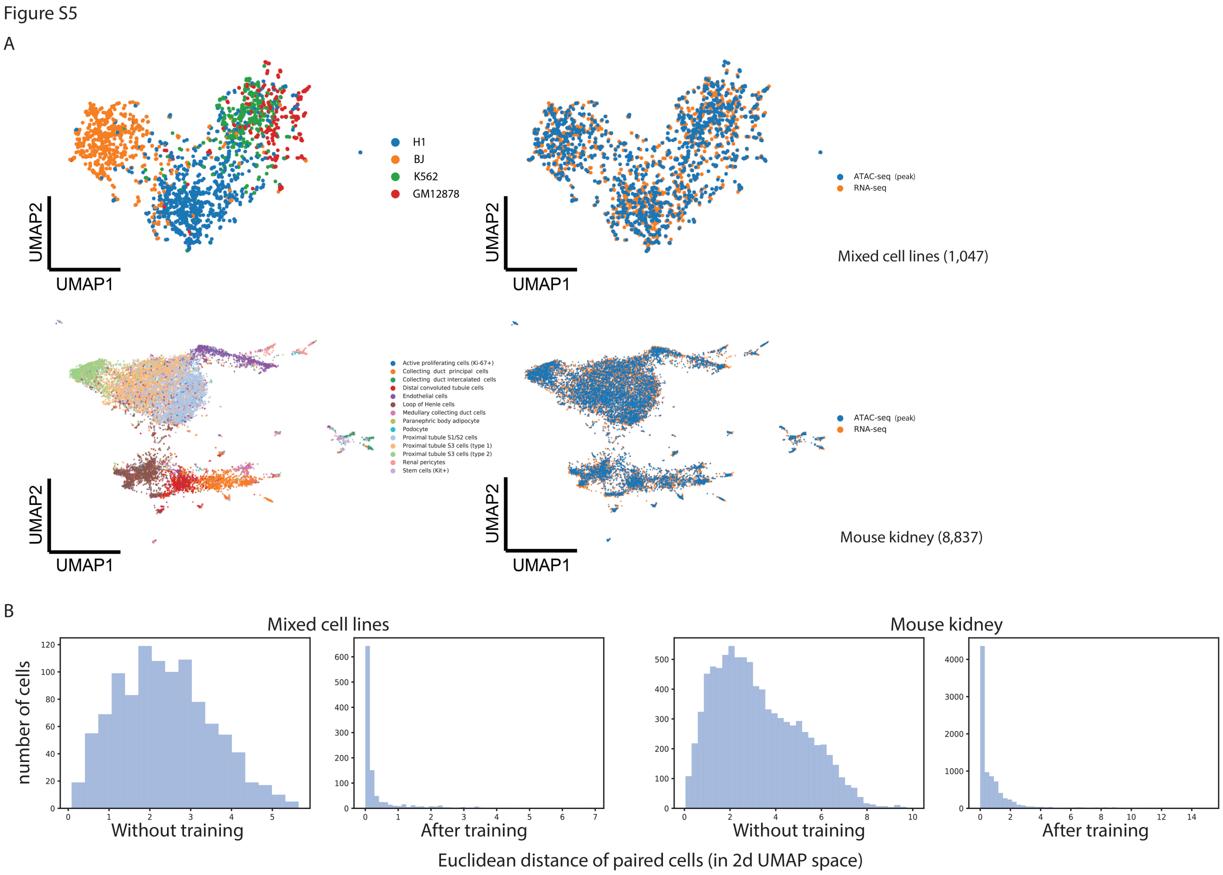


**Supplementary Fig. S5**

Integration of SNARE-seq and sci-CAR data through mpSMILE. a, UMAP visualization of integrated representation of mixed cell line data (upper) and mouse kidney data (lower) by mpSMILE. Cells are colored by cell-types (left) and data types (right). b, Histogram of Euclidean distance between paired cells, before and after training, based on the SMILE integrated 2D UMAP.

Figure S6

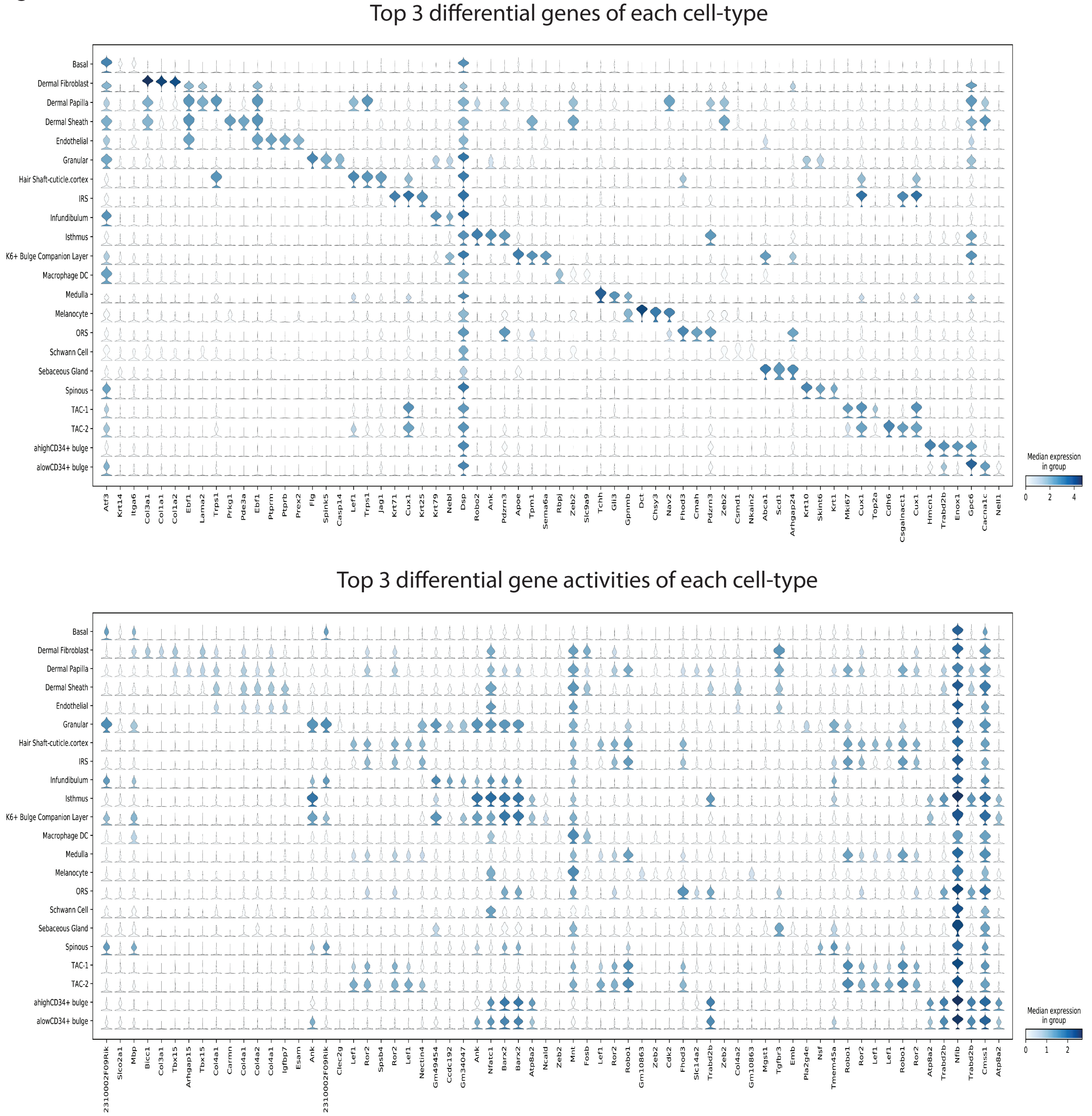


**Supplementary Fig. S6**

Differential gene expression and activity across 22 cell-types in mouse skin data. Only the top-3 genes for each cell-type were plotted.

Figure S7

A

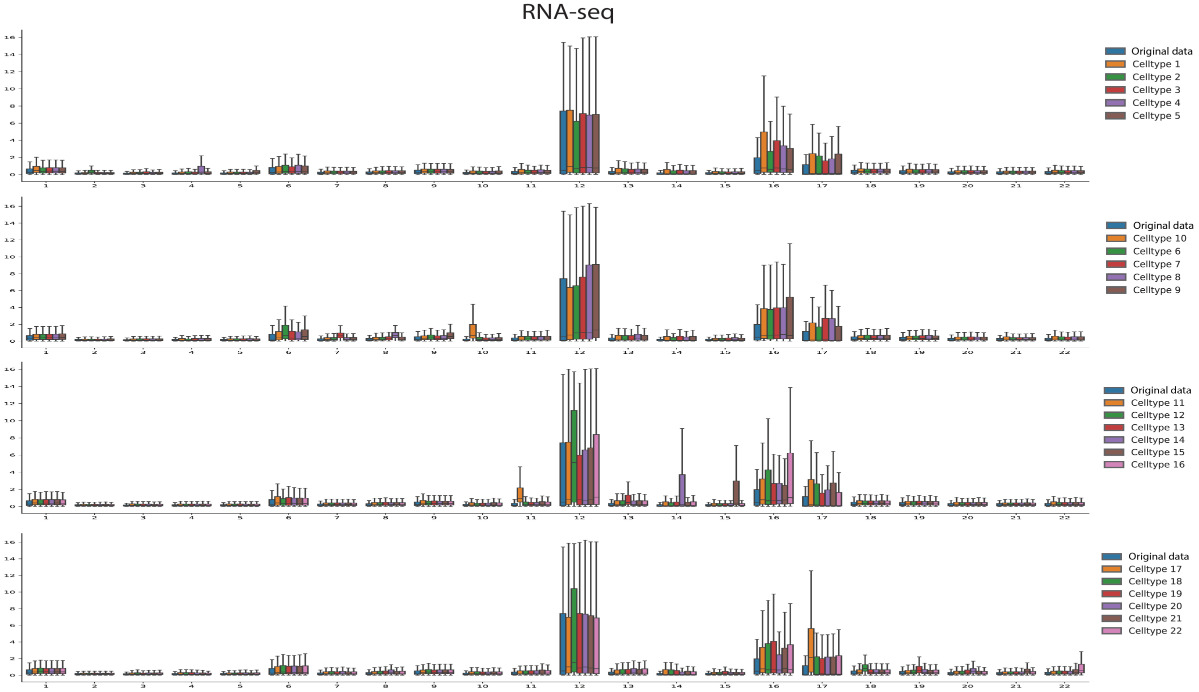


B

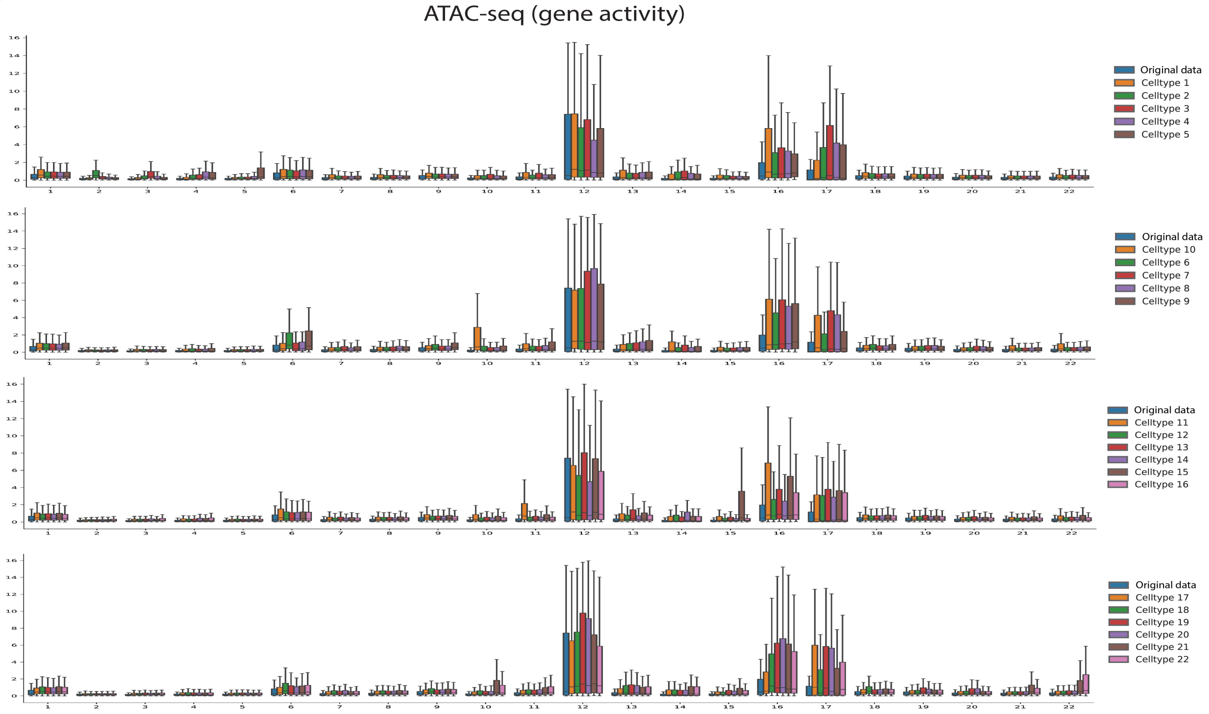


**Supplementary Fig. S7**

Explanation of co-embedding of RNA-seq and ATAC-seq by cluster-specific differential genes. a and b, boxplot of Euclidean distances of paired cells in 2D UMAP. Blue box: original data was forwarded through trained mpSMILE and Euclidean distances between cells in RNA-seq and their corresponding cells in ATAC-seq were measured in the integrated 2D UMAP. boxes of other colors: key differential genes that are specific to each cluster were suppressed to zeros, then the suppressed data were forwarded through trained mpSMILE, and Euclidean distances between cells in RNA-seq and their corresponding cells in ATAC-seq were measured in the integrated 2D UMAP. a) suppression of gene expression, and b) suppression of gene activity.

Figure S8

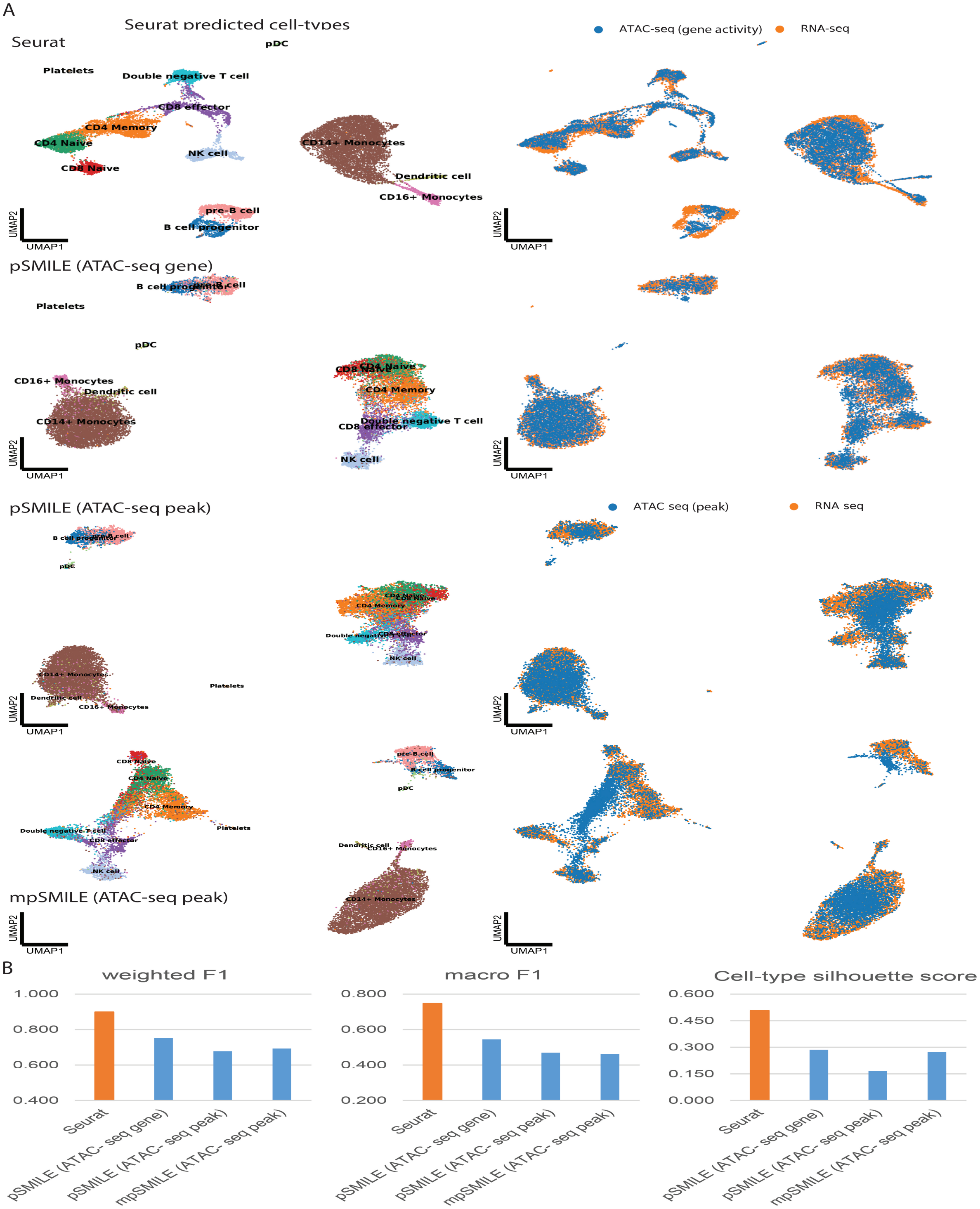


**Supplementary Fig. S8**

Integration of multimodal PBMC data through Seurat and pSMILE. a, UMAP visualization of integrated representation of PBMC data, by Seurat_v3 and p/mpSMILE with different ATAC-seq inputs. Cells are colored by cell-types predicted by Seurat_v3 (left panel) and colored by data types (right panel). b, Comparison of Seurat_v3 and p/mpSMILE in label transferring (RNA-seq to ATAC-seq) and data integration. Calculations of weighted F1, macro F1 and cell-type silhouette score are based on Seurat_v3 predicted cell-types as the ground truth.

Figure S9

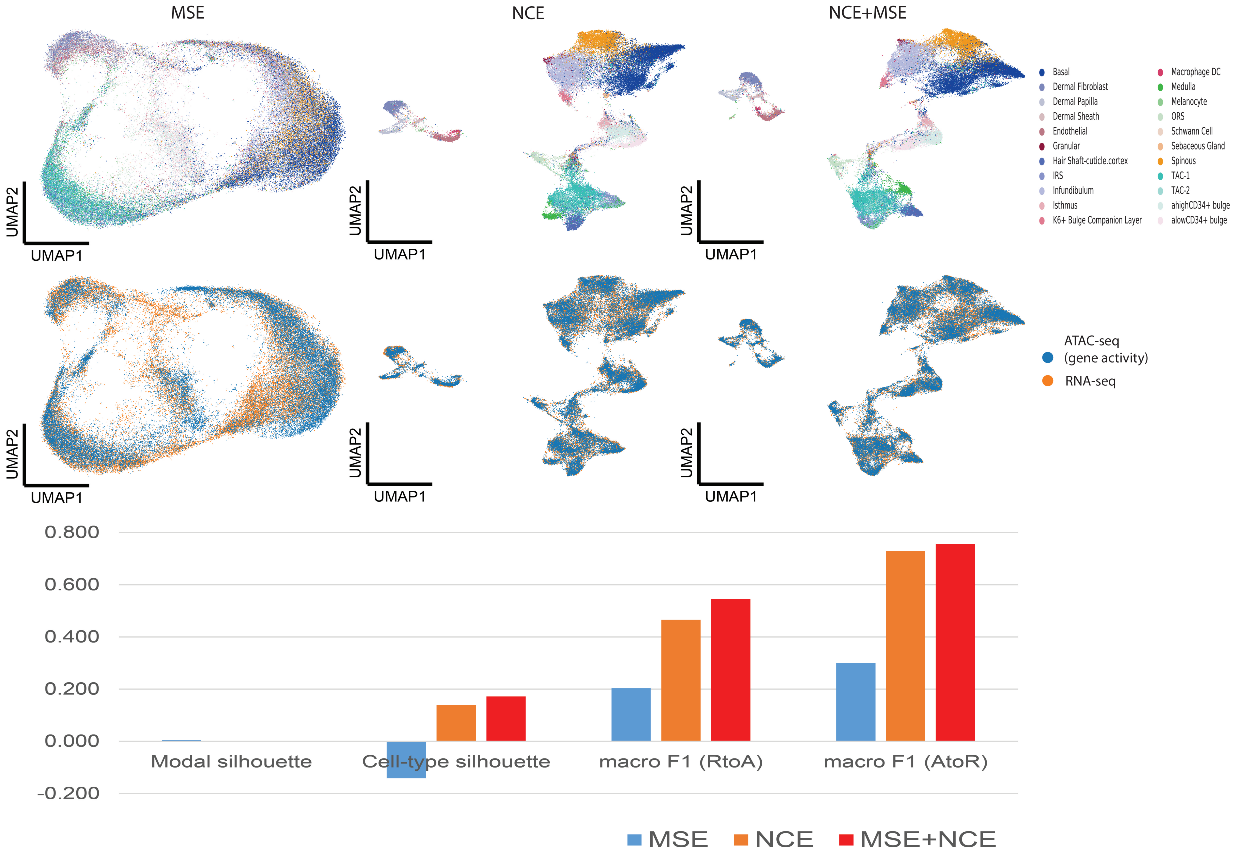


**Supplementary Fig. S9**

Integration and evaluation of mouse skin data through mpSMILE. mpSMILE was trained using MSE, NCE and NCE+MSE, respectively. Cells are colored by cell-types and data types. Modal and cell-type silhouette scores are based on data type and cell-type. label transferring from either RNA-seq to ATAC-seq or ATAC-seq to RNA-seq is evaluated through macro F1.

Figure S10

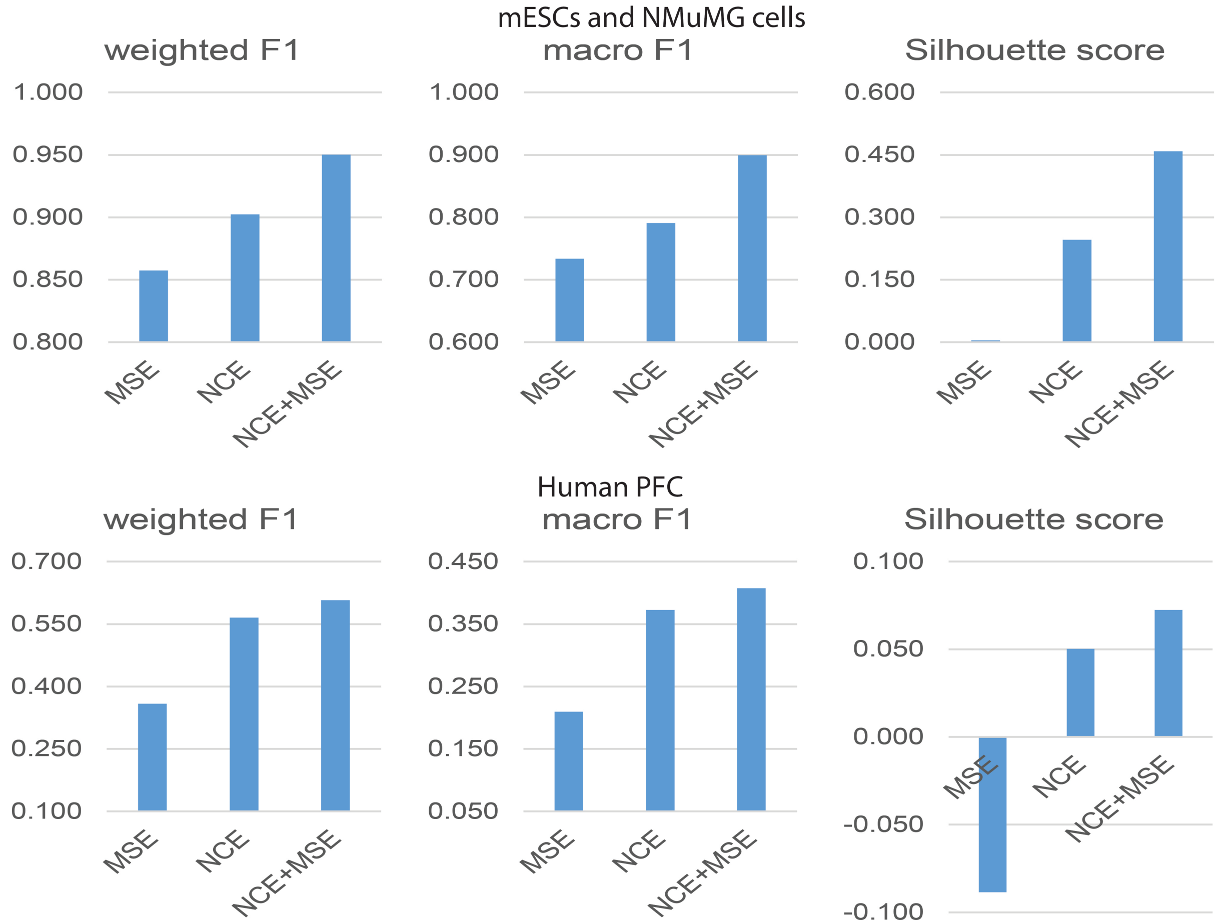


**Supplementary Fig. S10**

Evaluation of pSMILE with different loss functions. pSMILE was trained using MSE, NCE and NCE+MSE, respectively. Weighted F1 and macro F1 scores quantify label transferring from Methyl data to Hi-C data. Cell-type silhouette scores are based on author's cell-type label.
